# Sensitivity of *Listeria monocytogenes* to lysozyme predicts ability to proliferate in bovine caruncular epithelial cells

**DOI:** 10.1101/855841

**Authors:** Adam M. Blanchard, Rosemarie Billenness, Jessica Warren, Amy Glanvill, William Roden, Emma Drinkall, Grazieli Maboni, Robert S Robinson, Catherine E.D. Rees, Christiane Pfarrer, Sabine Tötemeyer

## Abstract

*Listeria monocytogenes* is an important foodborne pathogen in human and veterinary health, causing significant morbidity and mortality including abortion. It has a particular tropism for the gravid uterus, however, the route of infection in reproductive tissues of ruminants (i.e. placentome), is much less clear. In this study, we aimed to investigate a bovine caruncular epithelial cell (BCEC) line as a model for *L. monocytogenes* infection of the bovine reproductive tract. The BCEC infection model was used to assess the ability of 14 different *L. monocytogenes* isolates to infect these cells. Lysozyme sensitivity and bacterial survival in 580 µg lysozyme/ml correlated with attenuated ability to proliferate in BCEC (p=0.004 and p=0.02, respectively). Four isolates were significantly attenuated compared to the control strain 10403S. One of these strains (AR008) showed evidence of compromised cell wall leading to increased sensitivity to ß-lactam antibiotics, and another (7644) had compromised cell membrane integrity leading to increased sensitivity to cationic peptides. Whole genome sequencing followed by Multi Locus Sequence Type analysis identified that five invasive isolates had the same sequence type, ST59, despite originating from three different clinical conditions. Virulence gene analysis showed that the attenuated isolate LM4 was lacking two virulence genes (*uhpT*, *virR*) known to be involved in intracellular growth and virulence.

In conclusion, the BCEC model was able to differentiate between the infective potential of different isolates. Moreover, resistance to lysozyme correlated with the ability to invade and replicate within BCEC, suggesting co-selection for surviving challenging environments as the abomasum.

## Background

The zoonotic intracellular pathogen *Listeria monocytogenes* causes a range of clinical presentations including listeriosis, meningitis, septicaemia and abortions, in both cattle and humans. During pregnancy, *L. monocytogenes* is able to invade the placenta, causing inflammation leading to abortion by septicaemia [1, 2]. Listeriosis is of major veterinary importance in cattle due to its negative impact on animal health and the resulting economic losses [7].

The route by which *Listeria* spp. infect the ruminant placenta is unclear. Most studies have focused on infection of humans and rodents, and distinct species differences in placental structures as well as interhemal barriers mean that making comparisons between this and infection in other species is erroneous.

In the placenta, maternal and fetal tissues interact. In hemochorial placentas that are present in humans or guinea pigs, maternal blood comes into direct contact with fetal trophoblast cells. In contrast, in synepitheliochorial placentas found in cattle and sheep, the maternal and fetal blood are separated by several cell/tissue layers which any pathogen must cross to cause fetal infection [10]. In addition, the ruminant placenta is composed of multiple placentomes throughout the uterus, with each placentome formed from fetal cotyledons interdigitating with maternal caruncules. The latter are formed by multiple layers of stromal cells covered in a single layer of carunclar epithelial cells that interact with the fetal trophoblasts [10]. *L. monocytogenes* have been isolated from infected bovine placentomes post abortion and identified as the causative agent [11].

*Listeria* spp. invasion is primarily mediated by the interaction of the surface Internalin (Inl) proteins A and B with host cell receptors E-cadherin and c-Met tyrosine kinase (c-Met), respectively. For InlA-dependent entry into cells, proline at position 16 of E-cadherin is critical; in rats and mice if this proline is replaced by glutamic acid, then InlA-dependent entry into cells is prevented [12]. Whereas InlB-dependent cell invasion via c-Met does not occur in rabbits and guinea pigs but is functional in both mice and humans [13]. The *inlA* and *inlB* genes are arranged in an operon and can either be expressed as one bi-cistronic mRNA or independently expressed from promoters [14]. The overall pattern of gene expression is made more complex by the fact that there are multiple promoter sites which can be controlled by the virulence regulator, PrfA [14] and the stress sigma factor, Sigma B [15]. Generally, *inlA* mRNA levels are slightly higher than those for *inlB,* and expression of both genes is higher in the stationary phase of growth or under other environmental stress conditions [16]. However, it is also widely reported that many environmental strains may contain mutations in InlA which result in a less invasive phenotype [17], therefore, monitoring mRNA levels alone is not sufficient to fully characterise the virulence potential of strains.

Recently, InlP has been identified as a virulence factor linked to tissue tropism in the gravid uterus [18]. Deletion mutants of *inlP* were found to be attenuated using both, human explant and rodent models especially in the guinea pig, which most closely resembles the maternal-fetal interface of humans. The study suggested that InlP probably promotes pathogenesis at stages downstream of host cell invasion mediated by InlA or InlB (depending on the species), but there may be synergistic effects between InlP and InlA during infection of the placenta.

We hypothesize that L monocytogenes isolates from bovine abortions readily infect bovine caruncules and replicate within the cells. In this study, we investigated a bovine caruncular epithelial cell (BCEC) line as a model for *L. monocytogenes* infection of the bovine reproductive tract. The bovine E-cadherin and c-Met sequences and mRNA expression were analysed to determine permissiveness for interaction with InlA and InlB, respectively. The ability of a range of *L. monocytogenes* isolates from different clinical or environmental sources to infect the bovine caruncular epithelial cell lines was investigated. In addition, genome sequencing was used to determine MLST type, clonality and virulence gene presence of these isolates.

## Materials and Methods

### Bacterial culture

*Listeria monocytogenes* strains used in the infection studies are listed in Table 1. Bacteria were cultured overnight (approximately 17h) at 37°C in 5ml Heart infusion (HI) broth or on HI agar plates (Oxoid, UK). Growth was monitored using optical density (OD_600nm_) and cultures were diluted depending on the multiplicity of infection (MOI) required for infection experiments. The precise CFU/ml of the inoculum was then determined by serial dilution and plating on HI agar. To determine the growth rates and generation times of the isolates, overnight cultures were diluted in HI broth to OD600= 0.01. Growth was monitored using optical density (OD_600nm_) and serial dilution plated on HI agar.

**Table 1:** *Listeria monocytogenes* isolates used in this study.

| Strain number | Source<sup>a</sup> | PCR Serotype<sup>c</sup> | PCR Lineage<sup>d</sup> | MLST<sup>e</sup> | Generation time [min]<sup>i</sup> | Source/ Reference |
| --- | --- | --- | --- | --- | --- | --- |
| 10403S | Skin Lesion | 1/2a | II | 85 | 50±7.5 | (Bishop and Hinrichs, 1987) |
| AR008 | Healthy eye | 1/2a, 3a<sup>b</sup> | II<sup>b</sup> | 12 | 83.33±13.89<sup>f</sup> | (Warren et al., 2015) |
| C00938 | Kerato-conjunctivitis | 1/2a, 3a<sup>b</sup> | II<sup>b</sup> | 20 | 43.48±0.76 | APHA |
| R06262 | Kerato-conjunctivitis | 1/2b, 3b<sup>b</sup> | I<sup>b</sup> | 59 | 50±10 | APHA |
| C02118 | Kerato-conjunctivitis | 4b<sup>b</sup> | I<sup>b</sup> | 6 | 45.45±2.07 | APHA |
| LM7644 | Abortion | 1/2a, 3a<sup>b</sup> | II<sup>b</sup> | 122 | 62.5±11.72<sup>g</sup> | APHA |
| C08389 | Abortion | 1/2a, 3a<sup>b</sup> | II<sup>b</sup> | 7 | 58.82±17.3 | APHA |
| C08078 | Abortion | 1/2b, 3b | I | 59 | 52.63±5.54 | APHA |
| C07872 | Abortion | 1/2b, 3b | I | 59 | 55.56±6.17 | APHA |
| C04949 | Abortion | 1/2b, 3b | I | 59 | 38.46±8.88 | APHA |
| C07754 | Abortion | 1/2a, 3a | II | 91 | 38.46±2.96 | APHA |
| G03652 | Meningitis | 1/2b, 3b<sup>b</sup> | I<sup>b</sup> | 59 | 52.63±5.54 | APHA |
| LM4 | Milk | 1/2b, 3b<sup>b</sup> | I<sup>b</sup> | 1009 | 66.67±4.44<sup>h</sup> | (Lawrence et al., 1995) |
| LM6 | Milk | 4b<sup>b</sup> | I<sup>b</sup> | 1 | 52.63±5.54 | (Lawrence et al., 1995) |

APHA: Animal and Plant Health Agency <sup>a</sup> All isolates are from bovine sources except for the human isolate 10403S. <sup>b</sup> Warren et al., 2015 <sup>c</sup> Serotypes were determined using the PCR-based method of Doumith et al (2004). This method in conjunction with the lineage typing cannot distinguish between serotypes 1/2a and 3a or 1/2b and 3b. However, serotypes 3a and 3b are not commonly isolated <sup>d</sup> Lineages were determined using the PCR-based method of Ward et al. (2004) <sup>e</sup> Institute Pasteur *Listeria* MLST data base <sup>f</sup> Growth rate reduced compared to C04949 (p=0.001) <sup>g</sup> Growth rate reduced compared to C04949 (p=0.024) <sup>h</sup> Growth rate reduced compared to 10403S (p=0.039) <sup>i</sup> Growth rates presented as mean +/- SD (n=3)

### Multiplex PCR assay for *Listeria monocytogenes* serotyping

Multiplex PCR was performed in order to separate the four major serovars (1/2a, 1/2b, 1/2c, and 4b) and three main lineages (I, II, III) of *L. monocytogenes* [54, 55]. To prepare template DNA, three to six colonies resuspended in 1 ml of sterile water were incubated at 90°C for 10min and then chilled on ice for 10min; 1µl of this was used as template DNA for each PCR reaction.

### Cell culture and infections

Bovine caruncular epithelial cell line BCEC-1 (BCEC), provided by Prof. C. Pfarrer [56], were grown in DMEM/Ham’s nutrient mixture F12 1:1 (Sigma-Aldrich, UK) with 10% (v/v) fetal calf serum (Sigma), 2mM L-Glutamine and 100 U/ml penicillin/streptomycin (Gibco) at 37°C with 5% CO_2_ [57]. BCECs were seeded into 24-well plates (Thermo Scientific, UK) in 500µl of complete medium and grown to confluence. One hour before infection, the complete medium was replaced with antibiotic-free medium and the plate incubated at 37°C. Cells were infected with an MOI of 200 (n≥5, for details see result section), and incubated for 1h. Medium was then removed from the wells and replaced with medium containing 100µg/ml gentamycin (Sigma) to kill extracellular bacteria. After a further 1h incubation, the medium was replaced with medium containing 5µg/ml gentamycin and incubated for 2-24hr post-infection. To enumerate intracellular bacteria, cells were washed three times with pre-warmed (37°C) PBS and lysed by addition of 100µl of ice-cold 0.5% (v/v) Triton-X-100 (Fisher Scientific, UK) per well. This was incubated on ice for 20min and the resultant lysate serially diluted in PBS before 10µl samples were plated using the Miles Misra technique onto HI agar and incubated at 37°C overnight. Then, the CFU/ml of lysates was calculated.

### Antibiotic resistance screening

Samples (100µl) of each of the 14 isolates of *L. monocytogenes* were spread onto individual HI agar plates. Disks of penicillin G (1U), cefuroxime/sodium (30µg), oxacillin (1µg), ampicillin (25µg) and ciprofloxacin (1µg) (Oxoid Ltd, Basingstoke, UK) were immediately placed on top of the spread culture. The plates were then incubated at 37°C overnight and the zones of inhibition were measured (mm).

### Antimicrobial peptide challenge assay

The antimicrobial peptide challenge was performed as outlined by Burke *et al*. (2014) [35] using mouse cathelicidin-related antimicrobial peptide (H-GLLRKGGEKIGEKLKKIGQKIKNFFQKLVPQPEQ-OH; Isca Biochemicals, Exeter, UK) [58] at a final concentration of 10 µg/ml (stock concentration: 1mg/ml in dimethyl sulfoxide (DMSO)).

### Isolation of RNA, cDNA synthesis and quantitative (q) PCR

Late log phase culture containing approximately 10^9^ CFU/ml was centrifuged at 13000x*g* for 2 min at room temperature. The pelleted cells were suspended in 1ml RNAlater (Sigma Aldrich) and incubated for 1h at room temperature. The suspension was centrifuged at 13000x*g* for 5min and the supernatant removed. The pelleted cells were suspended in 375µl of freshly prepared cell wall disruption buffer (30 U/ml mutanolysin, 10mg/ml lysozyme in 10ml of 10mM Tris, 1mM EDTA buffer, pH 8), incubated at 37^°^C for 30 min and then centrifuged at 13000x*g* for 5min at room temperature. RNA was extracted using NuceleoSpin®RNA isolation kit (Macherey-Nagel, UK) following manufacturer’s instructions.

For BCEC RNA extractions, the supernatant was removed and cells were lysed with 350µl of RNA lysis buffer (Nucleospin®RNA isolation kits, Machery-Nagel, UK) followed by RNA isolation according to manufacturer’s instructions. Eluted RNA was quantified using Qubit (Qiagen) and stored at −80°C. RNA was diluted in water and cDNA was synthesized using MMLV reverse transcriptase (Promega, Madison, USA) according to manufacturer’s instructions. The final volume of each reaction was diluted in RNAse/DNAse free water (Fischer Scientific, UK).

Quantitative PCR was performed using a LightCycler® 480 (Roche, Hertfordshire, UK). For primer sequences see Table 2. For bacterial and host gene expression, qPCR was performed in 20µl reactions with 0.25mM of each of the forward and reverse primer, 2X Luminoct SYBR Green qPCR ready mix (Sigma-Aldrich, Dorset, UK), 25ng of cDNA and PCR grade water (Roche, Hertfordshire, UK). An initial denaturation cycle of 95°C for 10min was used followed by 45 cycles of 10s at 95°C, 50s at 60°C and 1min at 72°C and a final extension of 10min at 72°C. Normalized gene expression of each gene was calculated based on the method described by Hughes *et al* 2007 [59].

**Table 2:**
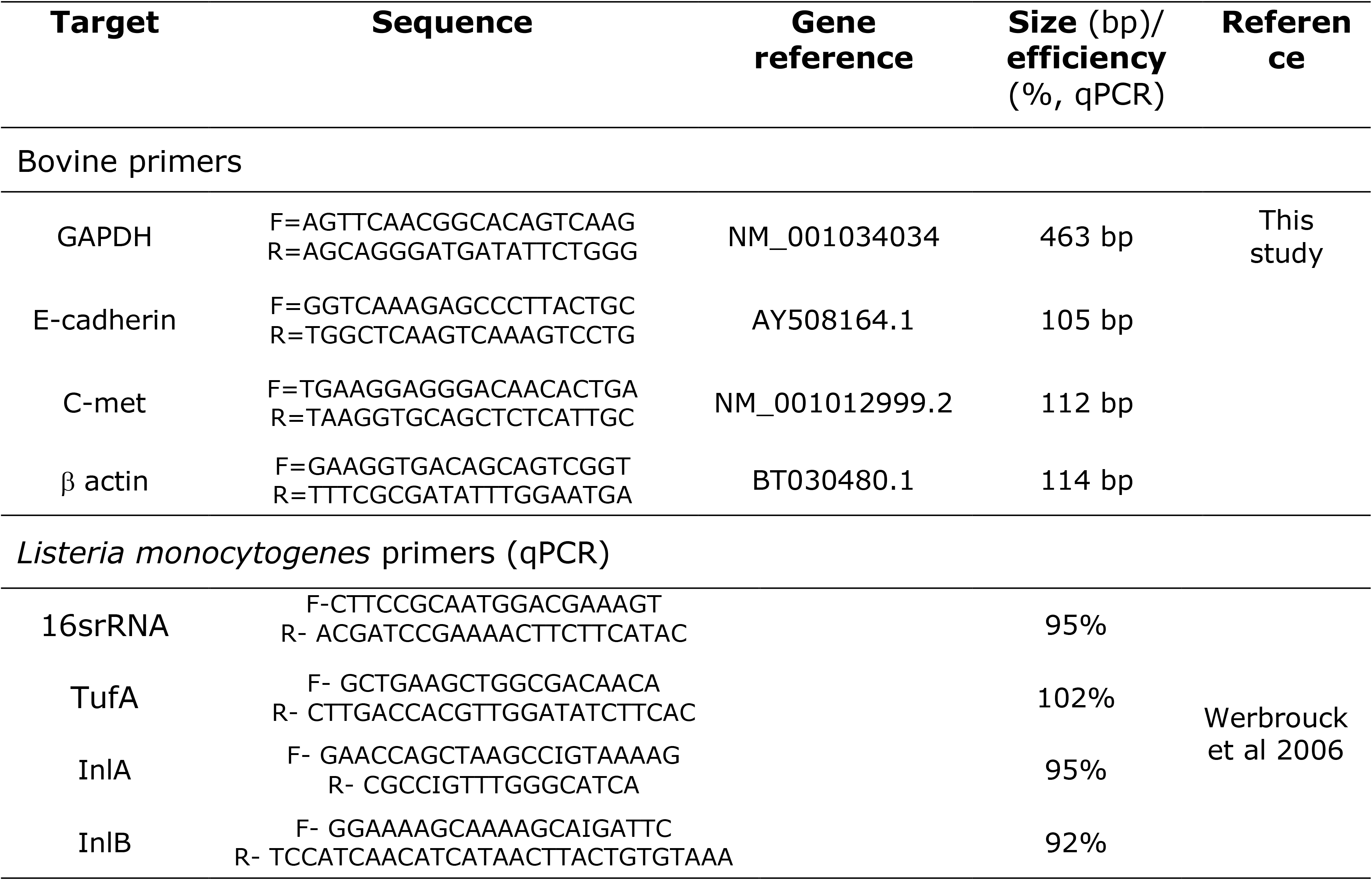
Primers.

### WGS and sequence analysis

DNA was extracted using the Cador Pathogen Minikit (Qiagen) following manufacturer’s recommendations. High throughput sequencing was performed at MicrobesNG (Birmingham U.K.) using Illumina MiSeq. Raw reads were assembled using the A5-MiSeq pipeline [60] and contigs were uploaded to the Pasteur MLST database were they are publically available and the MLST sequence type was determined (http://bigsdb.pasteur.fr/listeria/listeria.html).

### Multi Sequence Alignments

All nucleotide and protein alignments were completed using secondary structure aware high throughput multi-sequence alignment DECIPHER [61] (R script available at https://github.com/ADAC-UoN/DECIPHER-Sequence-Alignment.git). Trees were calculated using maximum likelihood by Fasttree double precision (version 2.1.8) [62] and visualised in iTOL [63].

### Virulence finder

Assembled genome files were uploaded to the virulence finder online *Listeria* database at the Danish Centre For Genomic Epidemiology (https://cge.cbs.dtu.dk/services/VirulenceFinder/version 1.5) [64] and searches were performed against reference isolate EDG-e, using a minimum of 90% identity along 80% of the coding sequence.

### Statistical Analysis

Statistical analysis of data was performed using GraphPad Prism 6.05. To compare the growth rates of *L. monocytogenes* isolates, a one-way ANOVA was carried out followed by Dunn’s multiple comparison test. To compare isolates in an infection context, a Kruskal-Wallis test was carried out, followed by Dunn’s multiple comparisons test. Pearson’s correlations were performed to compare data sets. *L. monocytogenes* sequence type distributions were analysed using Fisher’s exact test. Significance was reported for P<0.05.

## Results

### Sequence comparisons of host receptors E-cadherin and c-Met tyrosine kinase receptors

Since host specificity towards InlA-dependant entry into cells depends on the presence of proline at position 16 of E-cadherin in the first extracellular domain [12], alignment of the E-cad region of a range of species containing residue 16 was performed (Fig. 1A). This showed that bovine E-cadherin has proline at position 16 and suggests that bovine and ovine E-cadherin will interact with InlA in a similar way to human and guinea pig E-cadherin and can act as a receptor for *L. monocytogenes* in ruminant species. Interactions between InlB and c-Met are not as well defined; in c-MET the Sema, PSI and Ig1 region have shown to play a role in interaction with InlB [19]. Alignment of amino acids in these c-Met regions derived from bovine, ovine, human, murine, rabbit and guinea pig genome sequences showed that bovine c-Met does not cluster closely to rabbit and guinea pig c-Met (Fig. 1C). There were no consistent amino acid substitutions evident in the six amino acids of the Ig1 region that interacts with InlB (Fig. 1B), indicating that there is no obvious structural reason why InlB-dependent cell entry would not occur when *L. monocytogenes* interacts with bovine cells.

**Figure 1:**
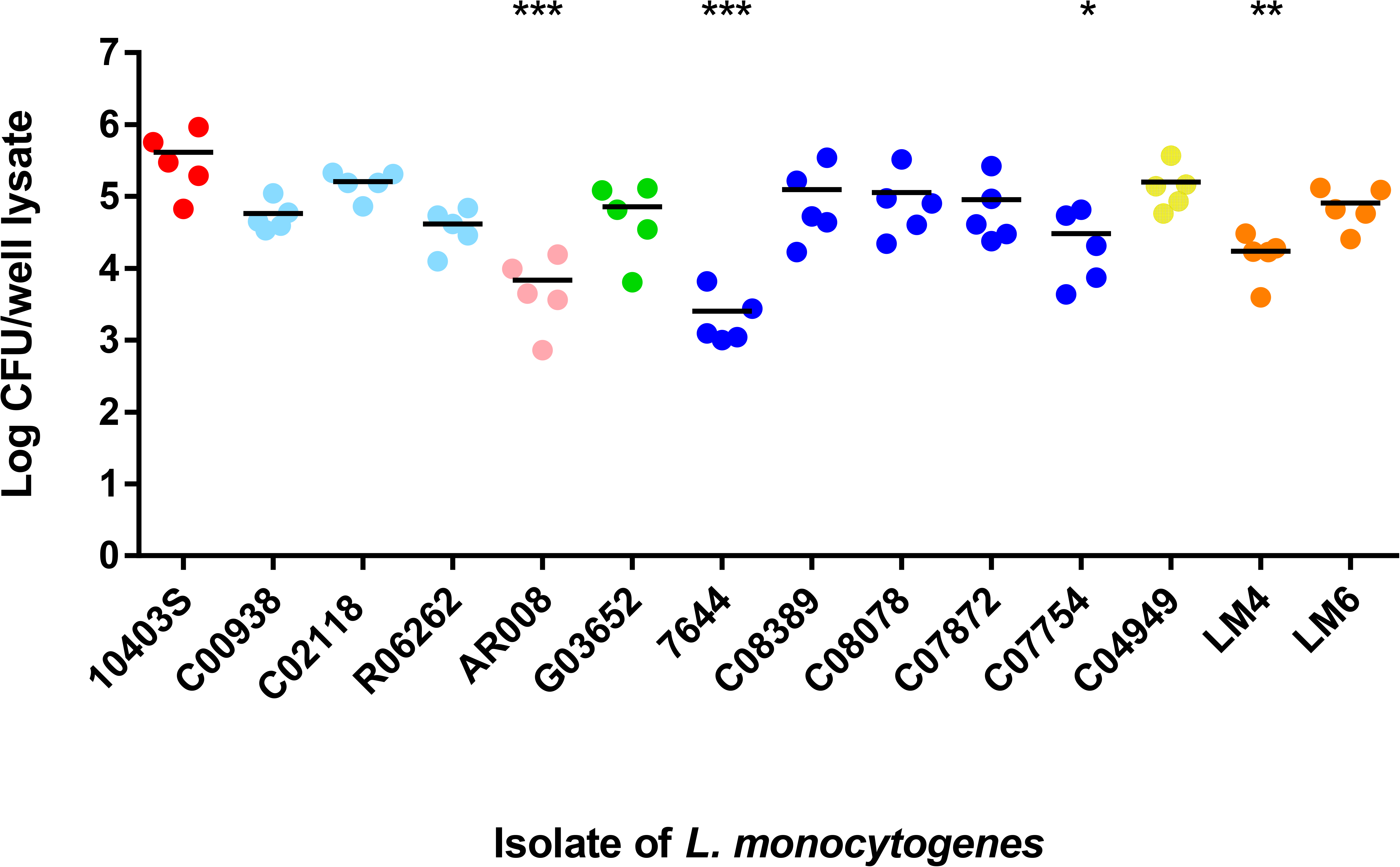
Bovine E-cad and cMet sequence comparisons and expression. (A) Multiple sequence alignment of the Pro16 residue of E-cad correlating with probability of invading the host cell [12], emboldened letters indicate amino acid substitutions previously identified in other studies and red letters indicate amino acid substitution unique to rodents. (B) Multiple sequence protein alignments of the region of c-MET, emboldened letters indicate amino acid substitutions which have previously been identified as a primary interface between InlB and c-MET [19], blue letters indicate relatedness of amino acid substitutions between certain species and red letters indicate amino acid substitutions discovered in this work. (C) Maximum likelihood tree of c-Met. E-cad (D) and c-MET (E) transcript levels in BCEC cells stimulated with 1µg.ml^-1^ LPS or infected with *L. monocytogenes* (MOI=200) for 4, 8 or 24 h.

Next, the expression of *E-cadherin* and *c-Met* mRNA in BCEC cells, chosen as the caruncular cell infection model was verified. mRNA for both were detected in these cells in the presence and absence of *L. monocytogenes* infection (Fig. 1D & E). More importantly, no difference in *E-cadherin* and *c-Met* mRNA expression level was observed when these cells were exposed to four different *L. monocytogenes* isolates and LPS (Fig. 1D & E). This indicated that infection with these bacteria did not down-regulate these receptors. Taken together, these results clearly demonstrate that *L. monocytogenes* should be able to productively interact with BCECs, and confirmed these cells are a suitable candidate to use as an infection model.

### *L. monocytogenes* infection of bovine caruncular epithelial cells

Initial experiments carried out to establish an infection method using the BCEC cells used a range of MOIs. This revealed that MOI of at least 200 was required to achieve consistent bacterial recovery from BCEC cells 2h post infection (data not shown), which is high but not unexpected as placental tissues are not easily or immediately invaded by *L. monocytogenes* [32]. Thus, all subsequent infections of BCEC cells were carried out using a MOI of 200. After 2h of infection, very low levels of intracellular bacteria (mean 0.78-1.5 log_10_ CFU per 2 x 10^5^ BCEC cells per well) were recovered and high levels of variability were observed between replicate infections. Although, the level detected was close to the detection limit (0.7 log_10_ CFU per 2 x 10^5^ BCEC cells per well) all isolates were able to invade BCEC 2h post infections to a similar extent (S1 Fig A). A preliminary time course of 4-24 h incubation post-infection showed that 24h yielded the most consistent and reproducible levels of bacterial recovery (S1 Fig B), therefore this was used for further experiments.

Using this infection model, the ability of the different *L. monocytogenes* isolates to invade BCEC cells was investigated. Fourteen *Listeria* spp. isolates from different origins and clinical presentations were used to infect BCEC cells including the well characterised strain 10403S (Table 1). Of the 14 isolates tested, four were significantly attenuated compared to the control strain (10403S). These were an isolate from a healthy bovine eye (AR008, P<0.001), an isolate from a milk processing plant (LM4 P<0.01) and two abortion isolates (7644 P<0.05, C07754 P<0.001) (Fig 2). In contrast to the control, the percentage of intracellular bacteria recovered was 1.9%, 8%, 0.5% and 33%, respectively (Fig. 2). No apparent extensive cell death was observed microscopically that would account for these low invasion rates.

**Figure 2:**
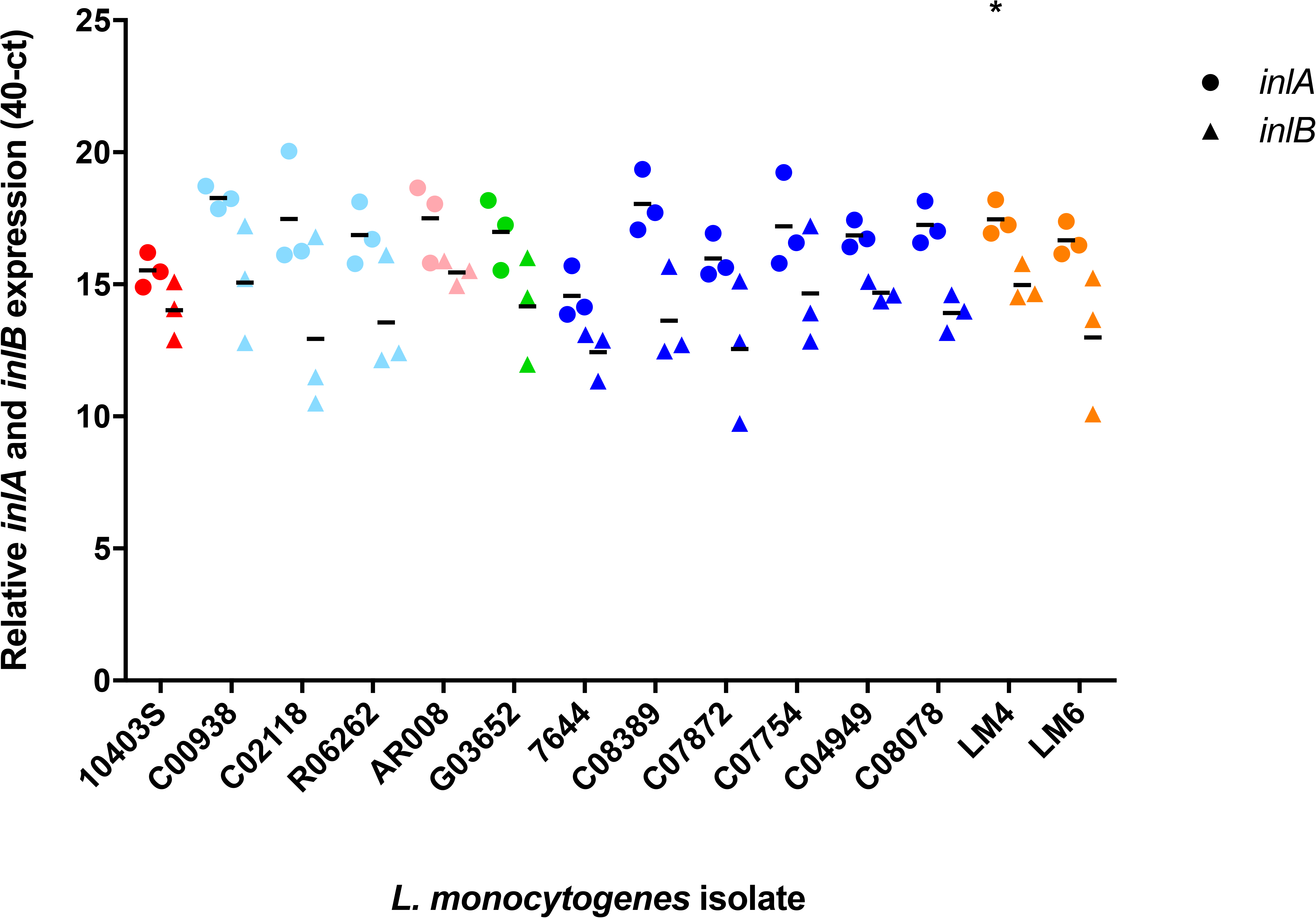
Infection of BCEC cells. BCEC cells were infected with an MOI=200 with *L. monocytogenes* isolates for 24 h at 37°C; For each isolate 5 independent experiments were performed. Dark blue indicates isolates from cases of bovine abortion, pale blue indicates isolates from bovine keratoconjunctivitis, orange indicates isolates from an environmental source, green indicates isolates from a case of meningitis, pink indicates an isolate from a healthy eye and red indicates the control strain 10403S originally a human isolate from a skin lesion. All data points and mean are shown. Statistical significance is shown compared to 10403S: *p<0.05, **p<0.01, ***p<0.001, (Kruskal-Wallis test followed by Dunn’s multiple comparisons test).

Differences in intracellular bacterial numbers may be due to the variation in InlA or B expression levels. Expression of *inlA* and *inlB* mRNA were determined in heart infusion broth as a proxy for the nutritional environment likely to be experienced in the animal host environment [20]. All isolates expressed *inlA* and *inlB* mRNA (Fig. 3) and this did not vary between strains (Fig. 3). As expected, *inlA* and *inlB* mRNA levels were positively correlated (r=0.58, p=0.03) and the levels of *inlB* mRNA were consistently lower than those of *inlA* as previously reported [21]. Thus, there was no evidence that differences in InlA or InlB levels would account for the differences in levels of intracellular bacteria recovered.

**Figure 3:**
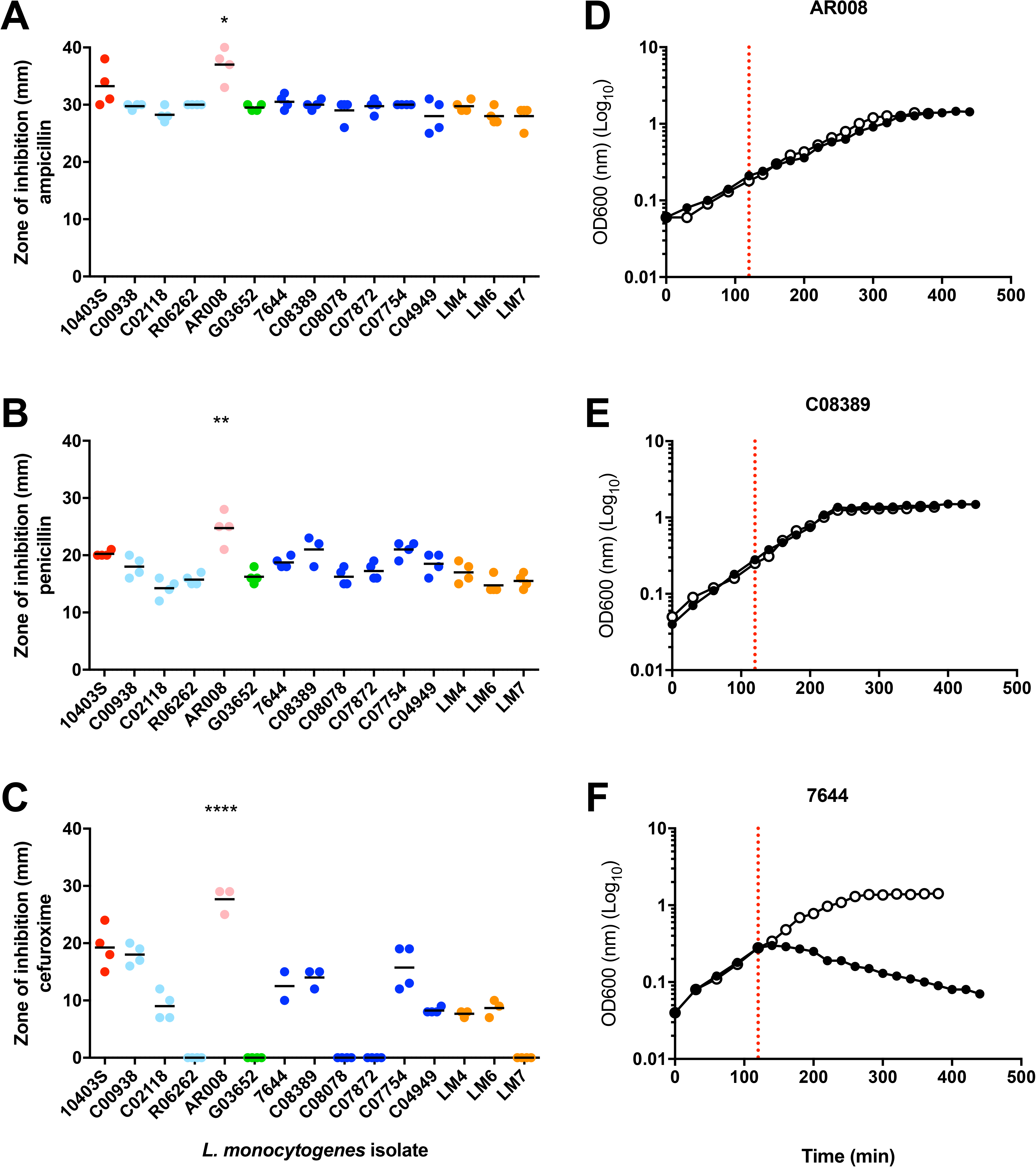
InlA and InlB expression. InlA and InlB transcript levels in *L. monocytogenes* isolates grown to late log phase in HI medium. Dark blue indicates isolates from cases of bovine abortion, pale blue indicates isolates from bovine keratoconjunctivitis, orange indicates isolates from an environmental source, green indicates isolates from a case of meningitis, pink indicates an isolate from a healthy eye and red indicates the control strain 10403S originally a human isolate from a skin lesion (Kruskal-Wallis test followed by Dunn’s multiple comparisons test).

### Sensitivity to lysozyme correlates with attenuated ability to proliferate in BCECs

Lysozyme sensitivity is an important factor in determining the ability of most *L. monocytogenes* strains to infect a bovine conjunctiva explant model [22]. Interestingly, the number of recovered bacteria from BCEC cells after 24h of infection correlated strongly with levels of lysozyme resistance (MIC [r=0.82, p=0.004]; bacterial survival in 580 µg lysozyme/ml [r=0.72, p=0.02]). However, there was no correlation with growth rate (r=0.49, p>0.05) or with intracellular bacteria recovered from conjunctiva explant infections (r=0.55, p>0.05) (Table 3, MIC & survival in lysozyme were previously reported) [22].

**Table 3:** Pearson correlations.

|  | **r** | **95% conf**<br>**interval** | **R<sup>2</sup>** | **p-**<br>**value** |
| --- | --- | --- | --- | --- |
| mRNA expression: *inlA* vs *InlB* | 0.58 | 0.07 to 0.85 | 0.34 | **0.03** |
| **Intracellular bacteria count (log**<br>**CFU/well with:**<br>**MIC lysozyme** | 0.82 | 0.40 to 0.96 | 0.67 | **0.004** |
| Survival in 580 µg/ml lysozyme (log<br>CFU/ml) | 0.72 | 0.16 to 0.93 | 0.52 | **0.02** |
| Growth rate in HI broth | 0.49 | -0.05 to 0.81 | 0.24 | 0.073 |
| Intracellular bacteria counts in<br>conjunctiva | 0.55 | -0.12 to 0.88 | 0.31 | 0.097 |

To investigate the basis of differences in sensitivity to lysozyme of these strains, isolates were challenged with β-lactam antibiotics (ampicillin, penicillin G and cefuroxime) to test cell wall integrity and a cationic peptide (mCRAMP) to test membrane integrity. Only one isolate, AR008 (isolated from a healthy eye), showed significant increased sensitivity compared to the reference strain 10403S to ampicillin (p=0.04), penicillin G (p=0.004) and cefuroxime (p=0.0001). This suggested that a compromised cell wall may contribute to the lysozyme sensitivity of this isolate (Fig. 4 A-C). Of the three lysozyme-sensitive isolates, only isolate 7644 showed sensitivity to mCRAMP, indicating that compromised cell membrane integrity may contribute to the lysozyme sensitivity of this isolate (Fig. 4 D-F).

**Figure 4:**
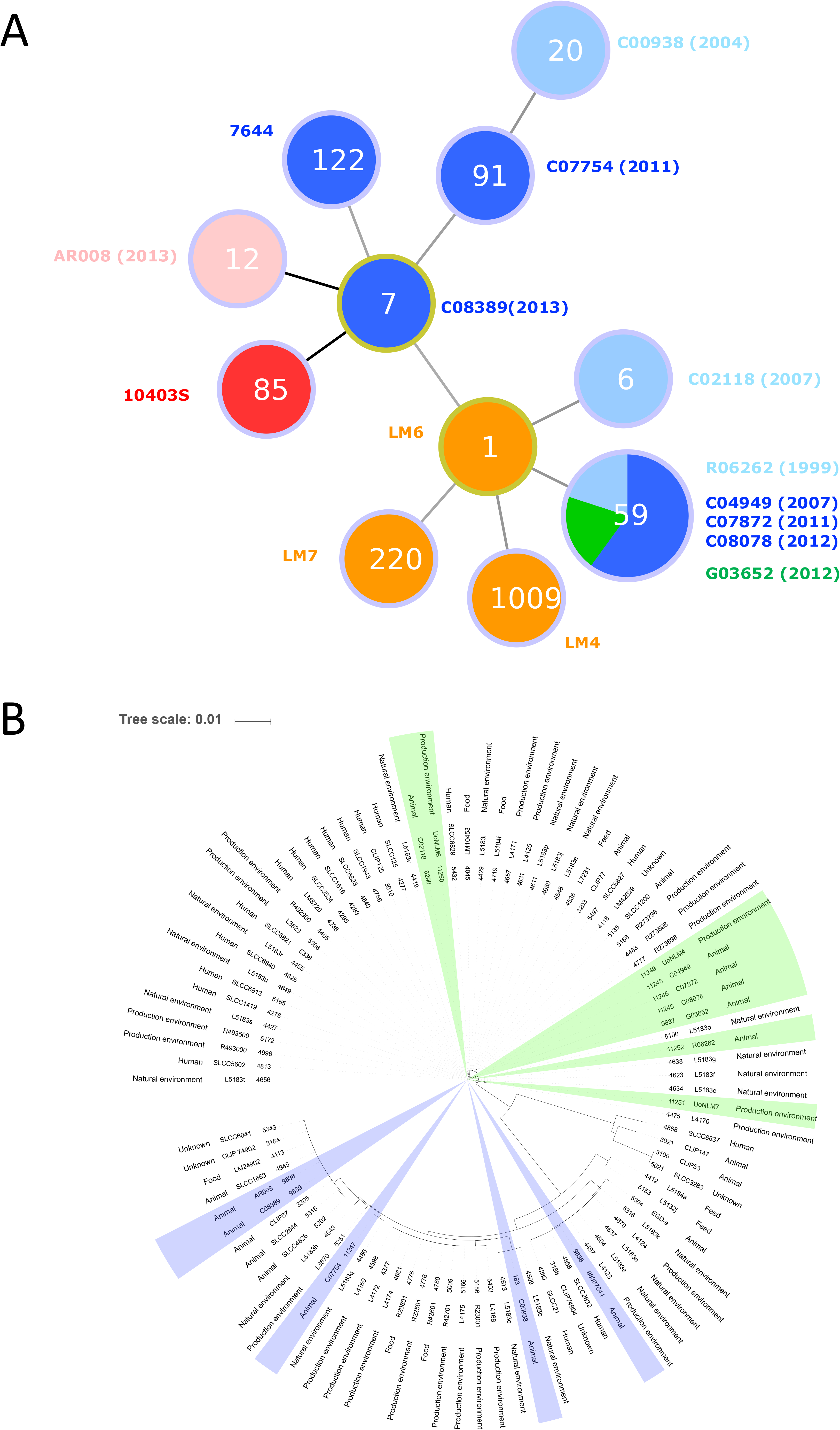
Treatment of *L. monocytogenes* with cell wall acting antibiotics and CRAMP. To investigate cell wall integrity, overnight cultures were plated on heart infusion agar and disks containing 1U penicillin G (A), 25μg ampicillin (B) or 30μg cefuroxime sodium (C). The plates were incubated overnight at 37°C, and zones of inhibition were measured. Dark blue indicates isolates from cases of bovine abortion, pale blue indicates isolates from bovine keratoconjunctivitis, orange indicates isolates from an environmental source, green indicates isolates from a case of meningitis, pink indicates an isolate from a healthy eye and red indicates the control strain 10403S originally a human isolate from a skin lesion. For each isolate 5 independent experiments were performed. All data points and mean are shown. Statistical significant increase in susceptibility is shown compared to 10403S using a. * P<0.05, ** P<0.01, *** P<0.001, **** P<0.0001. (One Way ANOVA followed by Dunnett’s multiple comparisons test). To assess cell membrane integrity, duplicate cultures of *L. monocytogenes* isolates AR008 (D), 7644 (E), C08389 (F) were grown to log phase in HI broth at 37°C and stimulated with a final concertation of 10mg.ml^-1^ CRAMP/DMSO or DMSO alone (red dashed line). Absorbance at 600 nm was measured in 20 min intervals. Data are representative of at least duplicate experiments.

### Analysis of *L. monocytogenes* sequence types, core genomes and virulence genes

To further characterise these isolates, WGS was carried out on all the uncharacterised isolates and these sequence data were added to the open access Pasteur MLST database to determine sequence types (ST) (Table 1, Fig. 5A). From the ten identified, eight were single STs, while five isolates belonged to ST59. Interestingly, while the ST59 isolates were collected over several years (1999-2012) and from three different clinical presentations (keratoconjunctivitis (n=1), meningitis (n=1), abortion (n=3)), they were all able to infect and replicate inside BCEC cells at levels comparable to wildtype (Fig. 5).

**Figure 5:**
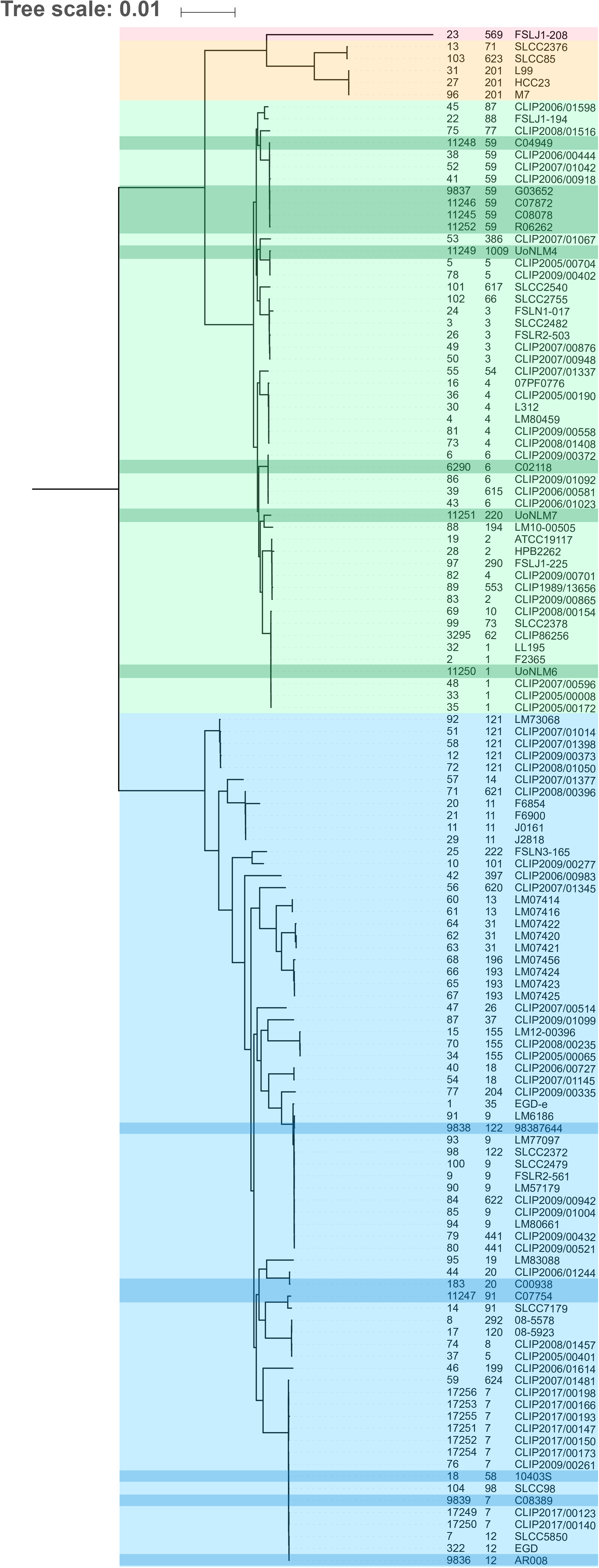
Epidemiological analysis of *L. monocytogenes* isolates based on MLST. (A) goeBURST analysis of the isolates used in this study; dark blue indicates isolates from cases of bovine abortion, pale blue indicates isolates from bovine keratoconjunctivitis, orange indicates isolates from an environmental source, green indicates isolates from a case of meningitis, pink indicates an isolate from a healthy eye and red indicates the control strain 10403S originally a human isolate from a skin lesion. Numbers in the circles denote the sequence type. (B) Maximum likelihood tree of the UK isolates present in the MLST database. Shaded areas correspond to the isolates used in this study, where green identifies linage I and blue identifies linage II.

Core genome analysis was carried out for the 128 isolates in the MLST database where a genome sequence was available (accessed 24/7/2017, S1 Table). This showed that isolates C00938 (ST20), C07754 (ST91), C02118 (ST6), AR008 (ST12) and LM6 (ST1) all cluster with other isolates of the same sequence type (Fig. 6). Some clusters contained more than one sequence type, for instance LM7644 (ST122) clustered with sequence types ST9, ST622 and ST441, and C08389 (ST7) is part of a cluster that also contains ST58 isolates (including 10403S) and ST98. Isolates LM4 (ST1009) and LM7 (ST220) were the only isolates of that sequence type present in the database but clustered closely with ST5 and ST194, respectively (Fig. 6).

**Figure 6:**
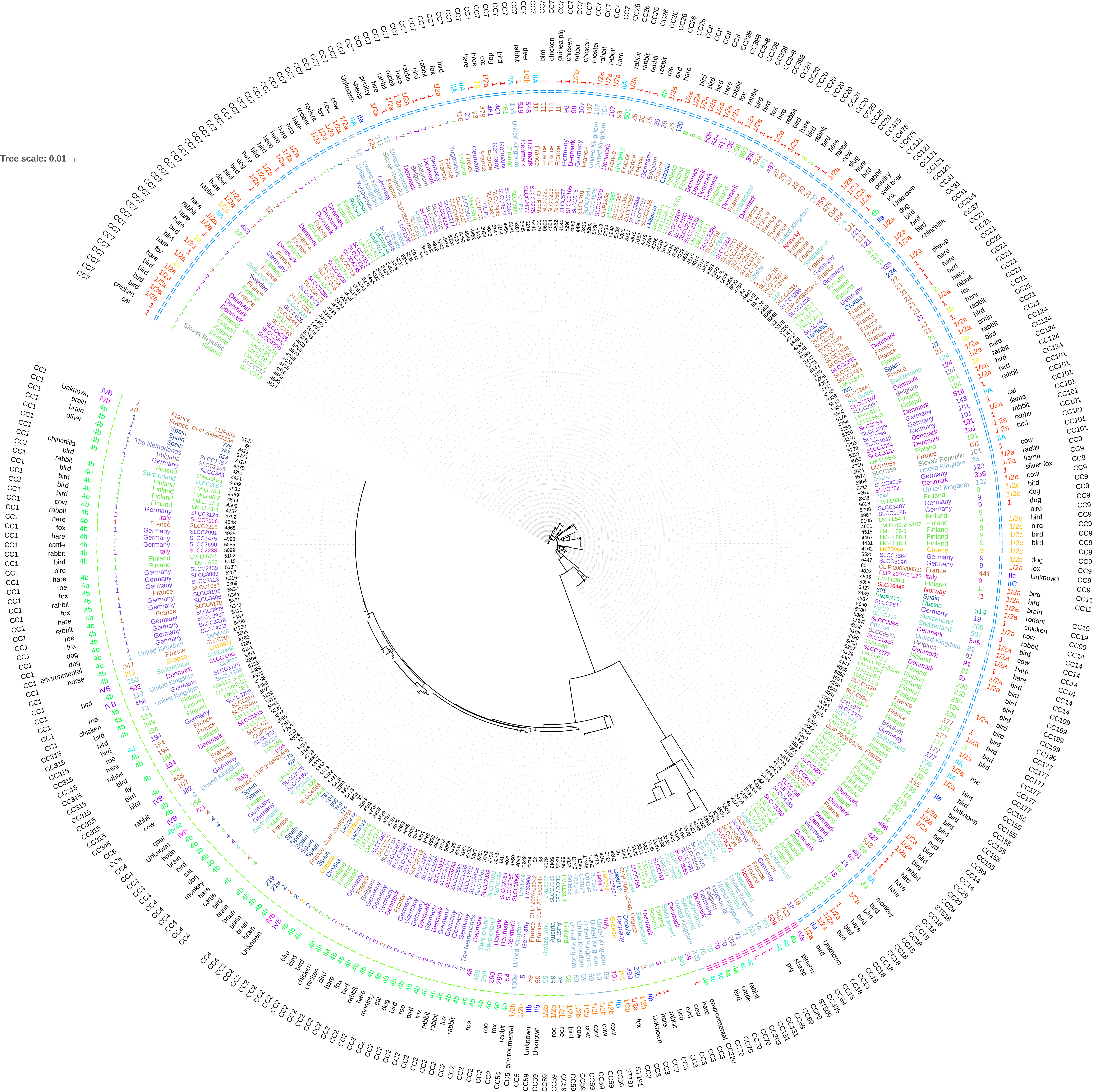
*L. monocytogenes* core genome comparison. Maximum likelihood tree has been generated using the core genome of isolates in the MLST database. Shaded areas correspond to linages, where green indicates linage I, blue indicates linage II, orange indicates linage III and red indicates linage IV. Darker shading highlights the isolates used in this study.

Further analysis of the WGS data for virulence gene content (presence and absence of genes as well as sequence similarity), showed that they clustered according to their linage and serotype as expected (Fig. 7, S2 Table). Sequence identity in general was high, between 91.3-100%, except for *inlK* (90.1-100%) and genes encoding the sRNA family *lhrC* (90.1-100 %) (Fig. 7, S2 Table). Five genes were only present in the six isolates of the 1.2a, 3a serotype: *inlL* (adherence), *ami* (adherence), *vip* (invasion) and two genes of the *lhrC* family (non-coding regulatory sRNA) (Fig. 7, S2 Table). The absence of those genes did not correlate with attenuation in the context of BCEC infection. In contrast, LM4 was the only isolate in this study that lacks two virulence genes known to be involved in intracellular growth and virulence. These were the *uhpT, the* sugar phosphate antiporter important for intracellular proliferation [23, 24] and *virR*, a transcriptional two component response regulator implicated in cell invasion and virulence *in vitro* and *in vivo* [25, 26] (S2 Table). In addition, LM4 had three other virulence genes (*srtB* (surface display), *sipZ* (intracellular survival) and *inlC* (internalin)) with the lowest reported level of sequence identity to the reference genome (S2 Table).

**Figure 7:**
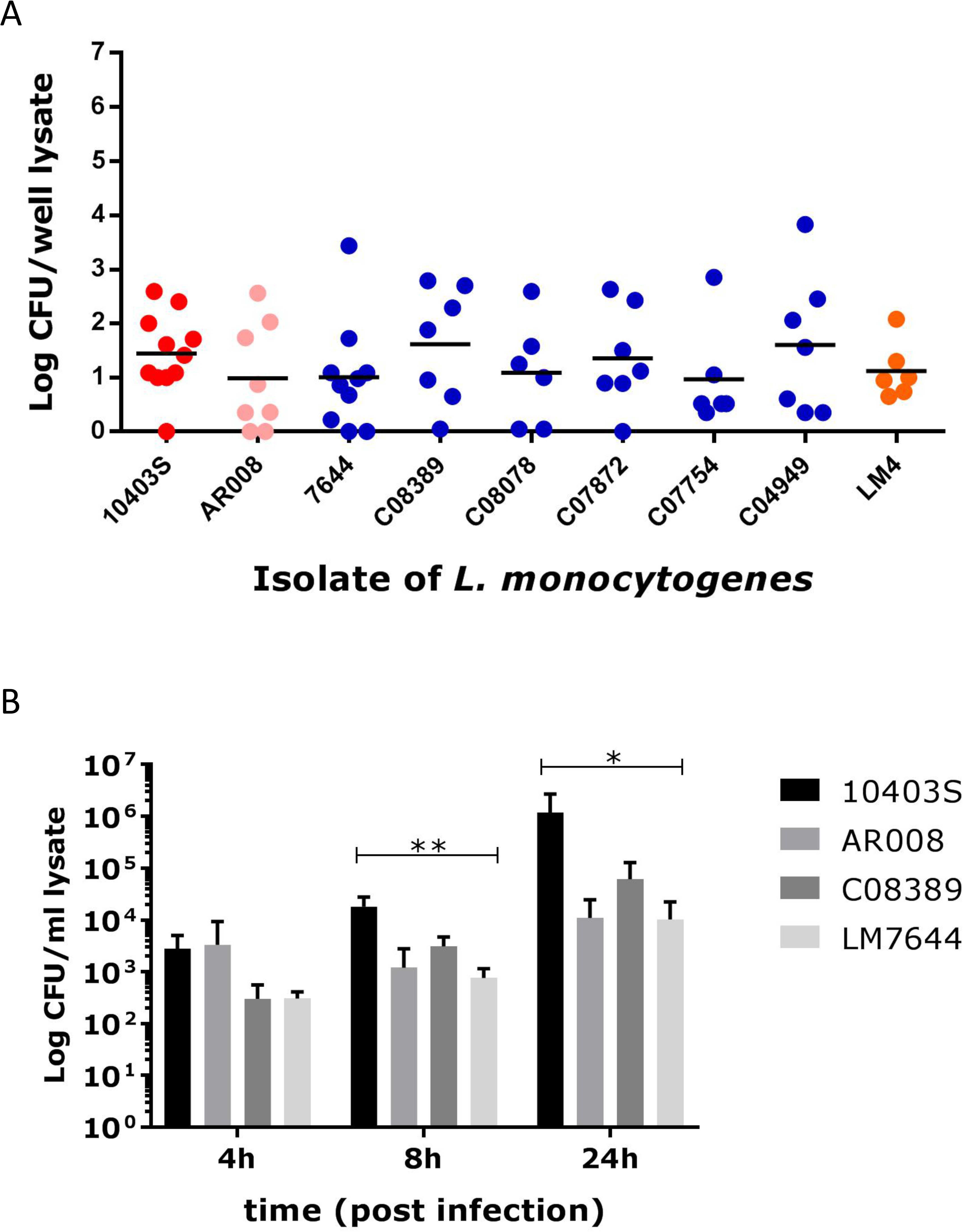
Virulence gene analysis. Heat map illustrating percentage identity of 87 *L. monocytogenes* virulence genes in comparison to isolate EDG-e determined through virulence finder [64], with crosses denoting the absence of genes. The gene matrix represents from top to bottom, genes involved in teichoic acid biosynthesis *(gtcA*), located in pathogenicity island LIPI-1 (*actA, hly, mpl, plcAB, prfA*), genes coding for internalins (*inlABCFHLKL*) and other genes involved in adherence (*ami, dltA, fbpA, lap, lapB*), invasion (*aut, iap, lpeA, recA, vip*), Intracellular survival (*clpBCEP, dal, fri, htrA, lplA1, oppA, perR, prsA2, pvcA, relA, sipZ, sod, svpA, tig, uHpt*), regulation of transcription and translation (*ctsR, fur, gmar, hfg*, *lhrc, lisKR, mogR, rsbv, sigB, stp, virk, rli55, rli60*), surface display (*lgt, lsp, sipX, srtAB, secA2*), peptidoglycan modification *(degU, murA, oatA, pgdA*), membrane integrity (*ctap, mrpf*), motility (*flaA, flgCE*), anaerobic growth (*eut*), regulation of metabolism (*codY*), immunomodulation (*chiA, lipA, lnyA, pgl*) and bile resistance (*bile, bsh*).

Analysis of the *inlA* sequences of these isolates for known changes that would be predicted to reduce levels of InlA (i.e. frameshifts causing premature stop codons or mutations in the promoter region [27]) was also performed. In agreement with the results gained from the *inlA* mRNA analysis (Fig. 3), no differences were identified in the *inlA* and *actA* promoter regions that might contribute to the lower levels of cell invasion and intracellular replication recorded. Similarly, InlP had 94% identity at the protein level across all of our isolates and therefore this did not seem to provide an explanation for the variation seen in the ability of these strains to infect the BCEC cells.

## Discussion

*L. monocytogenes* has been isolated from placentomes of infected cattle [1]. The maternal caruncle contains a dense network of blood vessels [28] allowing *Listeria* access to the maternal side of the placenta through the blood stream. The caruncle is also in close contact with the fetal chorion, meaning infection of the uterus can lead to endotoxaemia, an increased prostaglandin synthesis and subsequent lysis of the corpus luteum, leading to abortion. Alternatively, placentitis itself can disrupt the metabolic exchange of nutrients to the fetus, triggering the abortion [29]. While BCECs have been used for *L. monocytogenes* infections as a comparison to other tissues previously [30, 31] this study presents BCEC cells as infection model to characterise bovine *L. monocytogenes* isolates form different clinical presentations and sources. *L. monocytogenes* infected BCECs at low efficiency and required a high MOI. In other species, such as pregnant guinea pigs, colonisation of the placenta was initially slow with 10^3^-10^4^ fewer bacteria seen in the placenta than in the liver and spleen immediately after intravenous inoculation [32]. This suggests that placental tissues are not easily or immediately invaded by *L. monocytogenes*. This is consistent with our findings that low numbers of bacteria were recovered from caruncular cells 2h post infection with a wide range of variation. However, the invasion of a single bacterium into the placenta of guinea pigs can be sufficient to cause an abortion. Once colonised, there is poor bacterial clearance from the placenta and replication allows *Listeria* spp. to migrate into other tissues in high numbers [32]. This is consistent with our findings that at 24h post infection, higher numbers of bacteria were recovered from BCECs with less variation between infection experiments. This also suggests that any isolates able to invade BCECs may be able to cause an abortion *in utero*. Entry points towards systemic infections in cattle may include small breaches of oral mucosa from rough feeding material which may lead to repeat exposure of *L. monocytogenes* through contaminated silage [33].

The use of the BCEC infection model allowed us to identify strains with different potential to infect this cell type. Surprisingly, given that this isolate originated from a bovine abortion case, isolate 7644 was highly attenuated in the BCEC infection model. However, identification of *Listeria* from aborted fetuses is problematic, and the possibility exists that this isolate was a post-abortive environmental contaminant rather than the causative agent of infection. Alternatively, the animal may have been challenged with a high infectious dose. This assumption was previously proposed for a field isolate from a bovine abortion, which had a truncated PrfA, and was strongly attenuated in infection experiments with a wide range of cell types [31]. However, loss of infectivity may also be due to mutations accumulated during long term culture of the bacteria in a laboratory environment. In our previous study, both LM4 and 7644 were attenuated in a Caco2 infection model, whereas AR008 was able to infect Caco2 cells at similar levels to the control strain 10403S [22]. The fact that AR008 was attenuated in the BCEC model suggest that there are specific factors in these bovine placental cells involved in the interactions with *L. monocytogenes*.

Previously, we have shown that resistance to lysozyme was a positive predictive factor for infection of bovine conjunctiva explant model [22] but genome analysis performed in this study did not reveal any differences between the isolates used in the genes suspected to be involved in lysozyme resistance (*pdgA*, *oatA*, *degU*) [22, 34, 35]. Interestingly, there was also a strong correlation between *L. monocytogenes* replication in BCECs 24h post-infection and their level of lysozyme resistance. In cattle, lysozyme activity in most tissues is relatively low compared to other species, except for the abomasum [36] and tear fluid [37] which both have high levels of lysozyme activity. This may explain the co-selection for high levels of lysozyme resistance found in isolates from conjunctivitis as well as from other sites (reproductive tissues/fetus, brain and milk) that require the bacteria to survive passage through the abomasum. In addition, degradation of *L monocytogenes* cell wall by lysozyme leads to the release of peptidoglycan and its breakdown products that are ligands for the pattern recognition receptors of the innate immune system, such as Nod1, Nod2 and Toll-like receptor (TLR) 2 [38–41]. This is illustrated by *L monocytogenes* lacking *pgdA*, which was not only highly attenuated in its virulence *in vivo* and *in vitro* but also elicited a strong TLR2 and Nod1 dependent interferon-ß response [42]. This suggest that lysozyme resistance may contribute to *L monocytogenes* virulence in two different manners, by increasing bacterial survival as well as modulating the host response [43].

WGS identified the absence of virulence genes *virR* and *uhpT* in LM4 which potentially explains the attenuation of this isolate in the BCEC infection model. VirR is part of a two-component regulator (VirR–VirS) which is required for the virulence of *Listeria in vivo* [26]. It was also found that *virR* mutants are affected in their entry into Caco2 cells [25] and we have previously reported that LM4 also has a reduced capacity to invade this cell type [22]. The sugar phosphate antiporter UhpT promotes the uptake of phosphorylated hexoses during cytosolic growth [24] and deletion of this gene also leads to impaired intracellular proliferation in Caco2 cells [44]. Therefore, as LM4 lacks these two genes, it would be predicted that it would be less able to grow in BCEC cells. Interestingly, in our previous study, LM4 was not significantly attenuated in its ability to invade and proliferate in bovine conjunctiva tissues [22] but perhaps in that infection model the high level of resistance to lysozyme may compensate for any reduced intracellular growth. VirRS is also known to control the expression of a set of 17 genes, several of which affect bacterial cell wall and membrane integrity and, virR mutants are reported to be more sensitive to some beta-lactam antibiotics, including penicillin and cefuroxime [45]. However, LM4 did not show increased levels of sensitivity to these two antibiotics, or to challenge with cationic peptides, suggesting in the absence of VirR the genes in this operon are regulated in a different manner.

Genome sequencing of the isolates used in this study revealed that within our set of bovine clinical isolates, collected across the UK over several years and from different disease presentations, the MLST sequence type ST59 was over represented, with 5 out of the 10 clinical isolates belonging to that sequence type. Core MLST analysis further confirmed that they are closely related, forming a distinct cluster with other ST59 isolates. Within the MLST database, ST59 is associated with at least 6 human invasive infections (details are lacking for some human isolates) demonstrates that *L. monocytog*enes ST59 can be associated with invasive infections in both humans and cattle.

Interestingly, one of our cattle isolates has the MLST type ST6 (C02118, keratoconjunctivitis isolate, 2007), which is the sequence type identified in the large outbreaks of human disease in Europe and South Africa during 2017/2018. As sequence type and core genome analyses revealed that isolates from different clinical diseases, as well as from different species (human/cattle) cluster together, this suggests that it is less likely that the ability of *L. monocytogenes* to infect different host species is due to species-specific virulence factors, but more subtle variation in gene sequence influencing host interactions of this pathogen.

### Conclusion

*L. monocytogenes* is a highly versatile and adaptive bacterium, with the ability to not only infect a wide range of tissues within a host, but also to infect a wide range of physiologically distinct animal hosts. The placentome cell model provides a novel tool to characterise the infection processes carried out by *Listeria* spp. in a different host, where different host factors may influence the infection process.

## Declarations

### Ethics approval and consent to participate

Not Applicable

### Consent for publication

Not Applicable

### Availability of data and materials

The datasets analysed during the current study are available as part of the Pasteur institute MLST database repository, (http://bigsdb.pasteur.fr/listeria/)

### Authors’ contributions

ST, AMB and CEDR designed the experiments and wrote the manuscript, AMB, ST and RB analysed the data, RB, ED, JW, AG, WR and GM generated the data, CP provide cells lines and CP and RSR discussed experimental design. All authors have read and edited the manuscript.

## Acknowledgements

Not Applicable

## Additional information

**Additional File 1: Figure S1. Infection of BCEC cells for 2-24h.** BCEC cells were infected with an MOI=200 with *L. monocytogenes* isolates for 2-24 h at 37 °C. (A) BCEC cells were infected for 2 h with nine different isolates, for each isolate 6-11 independent experiments were performed. Dark blue indicates isolates from cases of bovine abortion, pale blue indicates isolates from bovine keratoconjunctivitis, orange indicates an isolate from an environmental source, pink indicates an isolate from a healthy eye and red indicates the control strain 10403S originally a human isolate from a skin lesion. All data points and mean are shown. (B) BCEC cells were infected for 4-24 h with four different isolates, for each isolate 3 independent experiments were performed, and average and standard deviation are shown. Statistical significance is shown: *p<0.05, **p<0.01.

**Additional File 3: Table S1. Multilocus sequence type metadata.** Metrics associated with all the isolates held in the MLST database.

**Additional File 4: Table S2. Determination of virulence associated genes.** Whole genome sequences were parsed through virulence finder [64] to identify putative virulence associated genes for the isolates used in this study. Green cells indicate greater than 99% similarity to virulent gene loci, orange cells indicate 95-98% similarity and red indicate between 90-94% similarity.

---

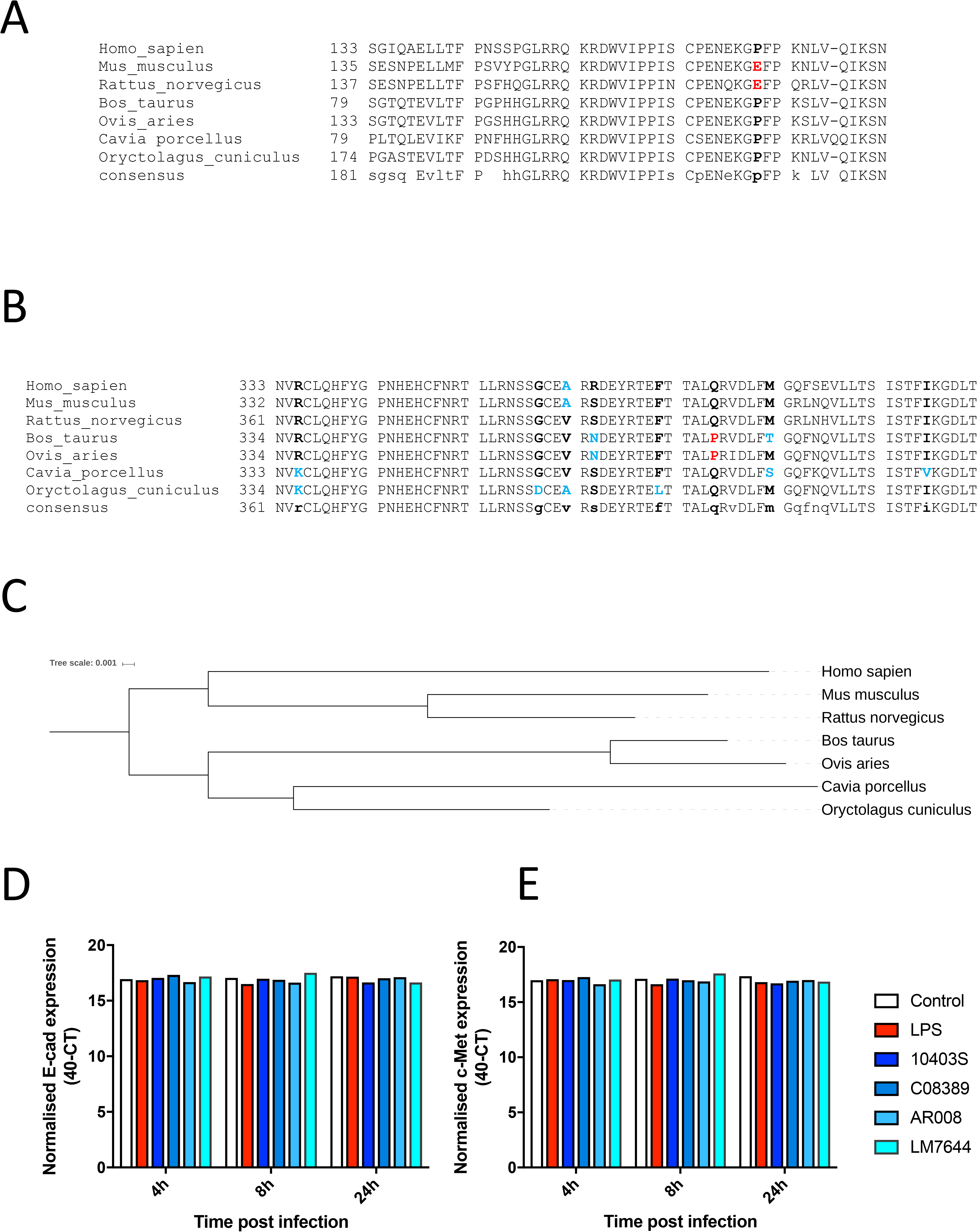

## References

1. Johnson CT, Lupson GR, Lawrence KE (1994) The bovine placentome in bacterial and mycotic abortions. Vet Rec 134:263 LP-266

2. Yaeger M, Holler L (2007) Bacterial causes of bovine infertility and abortion. In: Curr. Ther. large Anim. theriogenology. Saunders Elsevier, pp 389–399

3. Eurosurveillance editorial team (2016) The European Union summary report on trends and sources of zoonoses, zoonotic agents and food?borne outbreaks in?2015. EFSA J 14:20449

4. Department of Health Saouth Africa (2018) NICD Listeriosis Situation Report – 27 April 2018.

5. Food E, Authority S, Eurpoean Food Safety Authority (2018) Multi-country outbreak of Listeria monocytogenes serogroup IVb, multi-locus sequence type 6, infections probably linked to frozen corn. EFSA Support Publ. doi: 10.2903/sp.efsa.2018.EN-1402

6. Koopmans MM, Brouwer MC, Bijlsma MW, Bovenkerk S, Keijzers W, Van Der Ende A, Van De Beek D (2013) Listeria monocytogenes sequence type 6 and increased rate of unfavorable outcome in meningitis: Epidemiologic cohort study. Clin Infect Dis 57:247–253

7. Cabell E (2007) Bovine abortion: aetiology and investigations. In Pract 29:455–463

8. Erdogan HM, Cripps PJ, Morgan KL, Cetinkaya B, Green LE (2001) Prevalence, incidence, signs and treatment of clinical listeriosis in dairy cattle in England. Vet Rec 149:289–293

9. Animal and Plant Health Agency (2016) Veterinary Investigation Diagnosis Analysis (VIDA).

10. Robbins JR, Bakardjiev AI (2012) Pathogens and the placental fortress. Curr Opin Microbiol 15:36–43

11. Johnson CT, Lupson GR, Lawrence KE (1994) The Bovine Placentome in Bacterial and Mycotic Abortions. Vet. Rec.

12. Lecuit M, Dramsi S, Gottardi C, Fedor-Chaiken M, Gumbiner B, Cossart P (1999) A single amino acid in E-cadherin responsible for host specificity towards the human pathogen Listeria monocytogenes. EMBO J 18:3956– 3963

13. Khelef N, Lecuit M, Bierne H, Cossart P (2006) Species specificity of the Listeria monocytogenes InlB protein. Cell Microbiol 8:457–470

14. Lingnau A, Domann E, Hudel M, Bock M, Nichterlein T, Wehland JR, Chakraborty T (1995) Expression of the Listeria monocytogenes EGD inlA and inlB Genes, Whose Products Mediate Bacterial Entry into Tissue Culture Cell Lines, by PrfA-Dependent and -Independent Mechanisms. Infect Immun 63:3896–3903

15. Mcgann P, Wiedmann M, Boor KJ (2007) The Alternative Sigma Factor sigma B and the Virulence Gene Regulator PrfA Both Regulate Transcription of Listeria monocytogenes Internalins. Appl Environ Microbiol 73:2919–2930

16. Kim H, Marquis H, Boor KJ (2005) Sigma B contributes to Listeria monocytogenes invasion by controlling expression of inlA and inlB. Microbiology 151:3215–3222

17. Handa-Miya S, Kimura B, Takahashi H, Sato M, Ishikawa T, Igarashi K, Fujii T (2007) Nonsense-mutated inlA and prfA not widely distributed in Listeria monocytogenes isolates from ready-to-eat seafood products in Japan. Int J Food Microbiol 117:312–318

18. Faralla C, Rizzuto GA, Lowe DE, Kim B, Cooke C, Shiow LR, Bakardjiev AI (2016) InlP, a new virulence factor with strong placental tropism. Infect Immun 84:3584–3596

19. Niemann HH, Jäger V, Butler PJG, van den Heuvel J, Schmidt S, Ferraris D, Gherardi E, Heinz DW (2007) Structure of the Human Receptor Tyrosine Kinase Met in Complex with the Listeria Invasion Protein InlB. Cell 130:235– 246

20. Werbrouck H, Grijspeerdt K, Botteldoorn N, Van Pamel E, Rijpens N, Van Damme J, Uyttendaele M, Herman L, Van Coillie E (2006) Differential inlA and inlB expression and interaction with human intestinal and liver cells by Listeria monocytogenes strains of different origins. Appl Environ Microbiol 72:3862–3871

21. Tamburro M, Sammarco ML, Ammendolia MG, Fanelli I, Minelli F, Ripabelli G (2015) Evaluation of transcription levels of inlA, inlB, hly, bsh and prfA genes in Listeria monocytogenes strains using quantitative reverse-transcription PCR and ability of invasion into human CaCo-2 cells. FEMS Microbiol Lett 362:1–7

22. Warren J, Owen AR, Glanvill A, Francis A, Maboni G, Nova RJ, Wapenaar W, Rees C, Tötemeyer S (2015) A new bovine conjunctiva model shows that Listeria monocytogenes invasion is associated with lysozyme resistance. Vet Microbiol 179:76–81

23. Chico-Calero I, Suarez M, Gonzalez-Zorn B, Scortti M, Slaghuis J, Goebel W, Vazquez-Boland JA (2002) Hpt, a bacterial homolog of the microsomal glucose-6-phosphate translocase, mediates rapid intracellular proliferation in Listeria. Proc Natl Acad Sci 99:431–436

24. Chatterjee SS, Hossain H, Otten S, Kuenne C, Kuchmina K, Machata S, Domann E, Chakraborty T, Hain T (2006) Intracellular Gene Expression Profile of Listeria monocytogenes Intracellular Gene Expression Profile of Listeria monocytogenes †. Infect Immun 74:1323–1338

25. Mandin P, Fsihi H, Dussurget O, Vergassola M, Milohanic E, Toledo-Arana A, Lasa I, Johansson J, Cossart P (2005) VirR, a response regulator critical for Listeria monocytogenes virulence. Mol Microbiol 57:1367–1380

26. Camejo A, Buchrieser C, Couvé E, Carvalho F, Reis O, Ferreira P, Sousa S, Cossart P, Cabanes D (2009) In vivo transcriptional profiling of Listeria monocytogenes and mutagenesis identify new virulence factors involved in infection. PLoS Pathog. doi: 10.1371/journal.ppat.1000449

27. Moura A, Criscuolo A, Pouseele H, et al (2016) Whole genome-based population biology and epidemiological surveillance of Listeria monocytogenes. Nat Microbiol 2:16185

28. Senger P (2005) Pathways to Pregnancy and Parturition. Psychiatr Rehabil J 35:381

29. Miller RB (1977) A summary of some of the pathogenetic mechanisms involved in bovine abortion. Can Vet J 18:87–95

30. Rupp S, Bärtschi M, Frey J, Oevermann A (2017) Hyperinvasiveness and increased intercellular spread of listeria monocytogenes sequence type 1 are independent of listeriolysin s, internalin f and internalin J1. J Med Microbiol. doi: 10.1099/jmm.0.000529

31. Rupp S, Aguilar-Bultet L, Jagannathan V, Guldimann C, Drögemüller C, Pfarrer C, Vidondo B, Seuberlich T, Frey J, Oevermann A (2015) A naturally occurring prfA truncation in a Listeria monocytogenes field strain contributes to reduced replication and cell-to-cell spread. Vet Microbiol 179:91–101

32. Bakardjiev AI, Theriot JA, Portnoy DA (2006) Listeria monocytogenes traffics from maternal organs to the placenta and back. PLoS Pathog 2:0623–0631

33. Low JC, Donnelly W (1997) A Review of Listeria monocytogenes and Listeriosis. Veteriinaruy J 9–29

34. Aubry C, Corr SC, Wienerroither S, Goulard C, Jones R, Jamieson AM, Decker T, O’Neill LAJ, Dussurget O, Cossart P (2012) Both TLR2 and TRIF contribute to interferon-β production during listeria infection. PLoS One 7:1–9

35. Burke TP, Loukitcheva A, Zemansky J, Wheeler R, Boneca IG, Portnoy DA (2014) Listeria monocytogenes is resistant to lysozyme through the regulation, not the acquisition, of cell wall-modifying enzymes. J Bacteriol 196:3756–3767

36. Prieur DJ (1986) Tissue specific deficiency of lysozyme in ruminants. Comp Biochem Physiol -- Part B Biochem 85:349–353

37. Sotirov L, Semerdjiev V, Maslev T, Draganov B (2007) Breed-related differences in blood lysozyme concentration and complement activity in cows in Bulgaria. Rev Med Vet (Toulouse) 158:239–243

38. Kobayashi KS, Chamaillard M, Ogura Y, Henegariu O, Inohara N, Nunez G, Flavell RA (2005) Nod2-dependent regulation of innate and adaptive immunity in the intestinal tract. Science (80-). doi: 10.1126/science.1104911

39. Kobayashi K, Inohara N, Hernandez LD, Galán JE, Nüñez G, Janeway CA, Medzhitov R, Flavell RA (2002) RICK/Rip2/CARDIAK mediates signalling for receptors of the innate and adaptive immune systems. Nature. doi: 10.1038/416194a

40. Opitz B, Puschel A, Beermann W, Hocke AC, Forster S, Schmeck B, van Laak V, Chakraborty T, Suttorp N, Hippenstiel S (2006) Listeria monocytogenes Activated p38 MAPK and Induced IL-8 Secretion in a Nucleotide-Binding Oligomerization Domain 1-Dependent Manner in Endothelial Cells. J Immunol. doi: 10.4049/jimmunol.176.1.484

41. Torres D, Barrier M, Bihl F, Quesniaux VJ, Maillet I, Akira S, Ryffel B, Erard F (2004) Toll-like receptor 2 is required for optimal control of Listeria monocytogenes infection. Infect Immun

42. Boneca IG, Dussurget O, Cabanes D, et al (2007) A critical role for peptidoglycan N-deacetylation in Listeria evasion from the host innate immune system. Proc Natl Acad Sci U S A 104:997–1002

43. Ragland SA, Criss AK (2017) From bacterial killing to immune modulation: Recent insights into the functions of lysozyme. PLoS Pathog. doi: 10.1371/journal.ppat.1006512

44. Chico-Calero I, Suárez M, González-Zorn B, Scortti M, Slaghuis J, Goebel W, Vázquez-Boland JA (2002) Hpt, a bacterial homolog of the microsomal glucose-6-phosphate translocase, mediates rapid intracellular proliferation in Listeria. Proc Natl Acad Sci 99:431–436

45. Collins B, Curtis N, Cotter PD, Hill C, Ross RP (2010) The ABC transporter AnrAB contributes to the innate resistance of Listeria monocytogenes to nisin, bacitracin, and various β-lactam antibiotics. Antimicrob Agents Chemother 54:4416–4423

46. Wang Y, Zhao A, Zhu R, et al (2012) Genetic diversity and molecular typing of Listeria monocytogenes in China. BMC Microbiol 12:119

47. Huang YT, Ko WC, Chan YJ, Lu JJ, Tsai HY, Liao CH, Sheng WH, Teng LJ, Hsueh PR (2015) Disease burden of invasive listeriosis and molecular characterization of clinicalisolates in Taiwan, 2000-2013. PLoS One 10:4–15

48. Zilelidou EA, Rychli K, Manthou E, Ciolacu L, Wagner M, Skandamis PN (2015) Highly invasive Listeria monocytogenes strains have growth and invasion advantages in strain competition. PLoS One 10:1–17

49. Jennison A V., Masson JJ, Fang N-X, et al (2017) Analysis of the Listeria monocytogenes Population Structure among Isolates from 1931 to 2015 in Australia. Front Microbiol 8:1–13

50. Linke K, R??ckerl I, Brugger K, Karpiskova R, Walland J, Muri-Klinger S, Tichy A, Wagner M, Stessl B (2014) Reservoirs of Listeria species in three environmental ecosystems. Appl Environ Microbiol 80:5583–5592

51. Ebner R, Stephan R, Althaus D, Brisse S, Maury M, Tasara T (2015) Phenotypic and genotypic characteristics of Listeria monocytogenes strains isolated during 2011-2014 from different food matrices in Switzerland. Food Control 57:321–326

52. Dreyer M, Aguilar-Bultet L, Rupp S, et al (2016) Listeria monocytogenes sequence type 1 is predominant in ruminant rhombencephalitis. Sci Rep 6:36419

53. Jensen AK, Björkman JT, Ethelberg S, Kiil K, Kemp M, Nielsen EM (2016) Molecular Typing and Epidemiology. Emerg Infect Dis 22:625–633

54. Doumith M, Buchrieser C, Glaser P, Jacquet C, Martin P (2004) Differentiation of the Major Listeria monocytogenes Serovars by Multiplex PCR Differentiation of the Major Listeria monocytogenes Serovars by Multiplex PCR. J Clin Microbiol 42:3819–3822

55. Ward TJ, Gorski L, Borucki MK, Mandrell RE, Hutchins J, Pupedis K (2004) Intraspecific phylogeny and lineage group identification based on the prfA virulence gene cluster of Listeria monocytogenes. J Bacteriol 186:4994– 5002

56. Bridger PS, Haupt S, Klisch K, Leiser R, Tinneberg HR, Pfarrer C (2007) Validation of primary epitheloid cell cultures isolated from bovine placental caruncles and cotyledons. Theriogenology. doi: 10.1016/j.theriogenology.2007.05.046

57. Bridger PS, Menge C, Leiser R, Tinneberg HR, Pfarrer CD (2007) Bovine Caruncular Epithelial Cell Line (BCEC-1) Isolated from the Placenta Forms a Functional Epithelial Barrier in a Polarised Cell Culture Model. Placenta 28:1110–1117

58. Fritsche S, Knappe D, Berthold N, Von Buttlar H, Hoffmann R, Alber G (2012) Absence of in vitro innate immunomodulation by insect-derived short proline-rich antimicrobial peptides points to direct antibacterial action in vivo. J Pept Sci 18:599–608

59. Hughes S, Poh TY, Bumstead N, Kaiser P (2007) Re-evaluation of the chicken MIP family of chemokines and their receptors suggests that CCL5 is the prototypic MIP family chemokine, and that different species have developed different repertoires of both the CC chemokines and their receptors. Dev Comp Immunol 31:72–86

60. Coil D, Jospin G, Darling AE (2015) A5-miseq: An updated pipeline to assemble microbial genomes from Illumina MiSeq data. Bioinformatics 31:587–589

61. Wright ES (2015) DECIPHER: harnessing local sequence context to improve protein multiple sequence alignment. BMC Bioinformatics 16:322

62. Price MN, Dehal PS, Arkin AP (2010) FastTree 2 - Approximately maximum-likelihood trees for large alignments. PLoS One. doi: 10.1371/journal.pone.0009490

63. Letunic I, Bork P (2016) Interactive tree of life (iTOL) v3: an online tool for the display and annotation of phylogenetic and other trees. Nucleic Acids Res 44:W242–W245

64. Joensen KG, Scheutz F, Lund O, Hasman H, Kaas RS, Nielsen EM, Aarestrup FM (2014) Real-time whole-genome sequencing for routine typing, surveillance, and outbreak detection of verotoxigenic Escherichia coli. J Clin Microbiol 52:1501–1510

